# The biosynthetic gene cluster of the *C*-nucleoside antibiotic pyrazomycin with a rare pyrazole moiety

**DOI:** 10.1101/707406

**Authors:** Guiyun Zhao, Shunyu Yao, Kristina W Rothchild, Tengfei Liu, Yu Liu, Jiazhang Lian, Hai-Yan He, Katherine S Ryan, Yi-Ling Du

## Abstract

Pyrazomycin is a rare *C*-nucleoside antibiotic with a naturally occurring pyrazole ring, whose biosynthetic origin has remained obscure for decades. In this study, we report the identification of the gene cluster responsible for pyrazomycin biosynthesis in *Streptomyces candidus* NRRL 3601, revealing that StrR-family regulator PyrR is the cluster-situated transcriptional activator governing pyrazomycin biosynthesis. Furthermore, our results from *in vivo* reconstitution and stable-isotope feeding experiments support that PyrN is a new nitrogen-nitrogen bond forming enzyme linking the ε-NH_2_ nitrogen of l-*N*^6^-OH-lysine and α-NH_2_ nitrogen of l-glutamate. This study lays the foundation for further genetic and biochemical characterization of pyrazomycin pathway enzymes constructing the characteristic pyrazole ring.

## Main text

Nitrogen-nitrogen (N-N) containing natural products are group of specialized metabolites with diverse structures and a variety of biological activities.^[1]^ These molecules have been isolated from different sources, including bacteria, fungi and plants. Despite extensive studies on the genetic and biochemical basis of natural product biosynthesis over the past three decades, the biochemical routes leading to enzymatic N-N bond formation are only starting to be revealed.^[2–14]^ We recently reported a heme-dependent piperazate synthase responsible for N-N cyclization in piperazic acid, a building block for many nonribosomal peptide (NRP) or NRP-polyketide hybrid molecules (**Fig. 1**).^[2]^ The biosynthetic route to piperazate begins with the *N*-hydroxylation of l-ornithine, giving a hydroxylamine precursor, which is reminiscent of valanimycin biosynthesis, which also begins with a hydroxylamine precursor.^[9]^ By contrast, N-N bond formation in other N-N bond containing natural products including compounds cremeomycin, fosfazinomycin and kinamycin starts instead with the generation of nitrous acid,^[5,6,8,10]^ and the biosynthesis of the *N*-nitroso streptozocin originates with oxidation of the guanidine of l-arginine.^[3,4]^ However, whether other pathways to N-N bond containing molecules might begin from hydroxylamine precursors was unknown. Recent studies into hydrazone unit formation in the dipeptide s56-p1 resulted in the discovery of a pathway that links the hydroxylamine l-*N*^6^-OH-Lys to hydrazino acetic acid formation.^[7]^ In this pathway, Spb40, a fusion protein with cupin and aminoacyl-tRNA synthetase (aaRS)-like domains, appears from expression studies in *E. coli* to couple glycine and l-*N*^6^-OH-Lys by forming a N-N bond between the α-NH_2_ of glycine and the ε-N atom of l-*N*^6^-OH-Lys, the latter of which is in turn generated by Spb38-catalyzed *N*^6^-hydroxylation of l-Lys. An Spb40-like enzyme was also reported in the gene cluster for triacsins, but the details of this reaction, and whether this type of enzymology might play a role in pathways to other N-N bond containing structures, remains obscure.^[14]^

**Figure 1.**
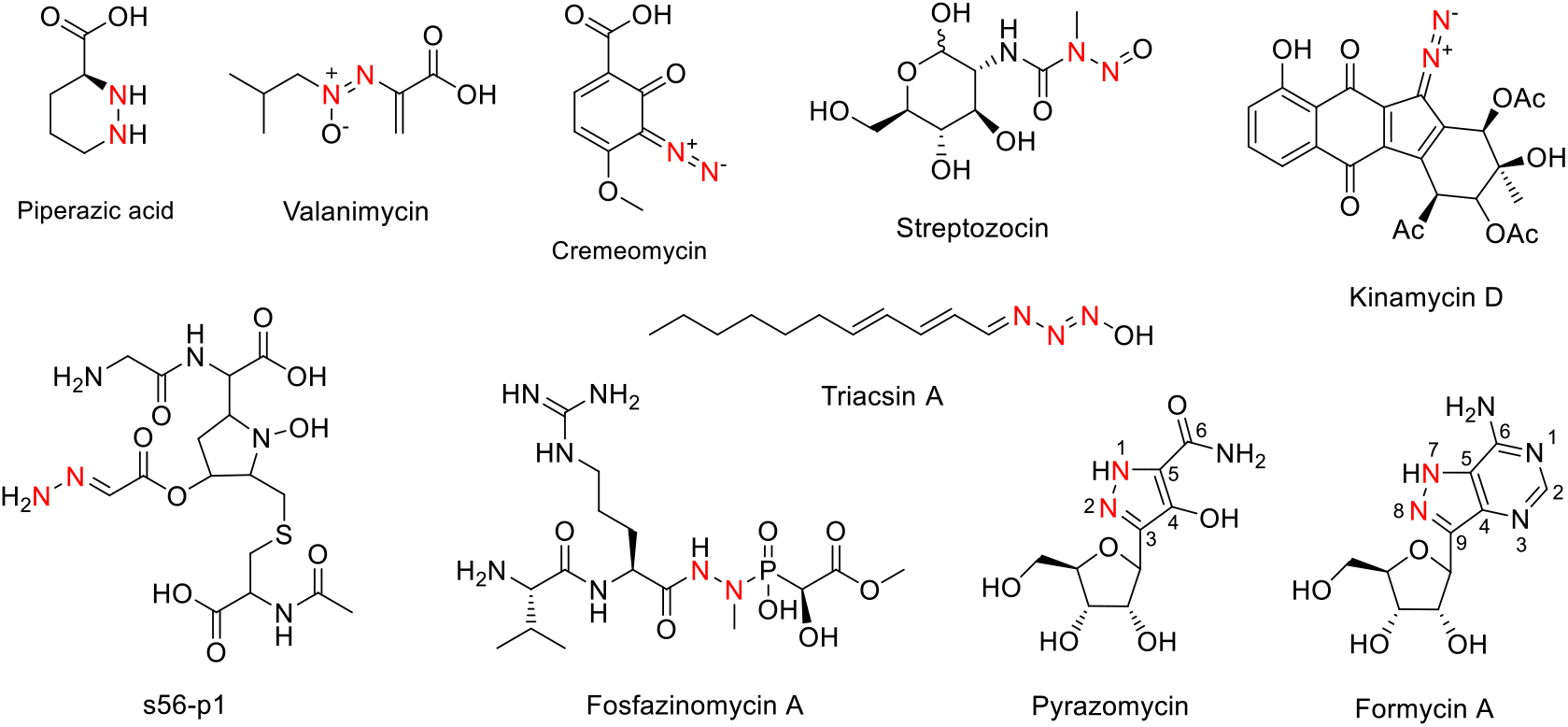
Examples of natural products containing a nitrogen-nitrogen linkage.

Despite progress toward understanding the enzymology of N-N bond formation, biosynthetic routes to aromatic structures containing N-N bonds is largely unexplored. Construction of an N-N bond embedded in an aromatic ring might require a very different biosynthetic logic from formation of a hydrazine or other N-N linkage. Among N-N bond-containing structures, a key example is pyrazomycin (PZN, also known as pyrazofurin), a *C*-nucleoside with a rare pyrazole moiety.^[15]^ As a nucleoside analog, PZN is a potent inhibitor of orotidine 5′-monophosphate decarboxylase and possesses broad-spectrum antiviral and antitumor activities.^[16]^ Early biosynthetic studies on PZN and its structural analog formycin used isotope-labeled precursors to establish that the C-3 to C-6 of the pyrazole ring in PZN derive from C-4 to C-1 of glutamate or α-ketoglutarate. Similarly, the C-9, C-4, C-5, C-6 of the pyrazolopyrimidine ring in formycin derive from these precursors (**Fig. 1**).^[17]^ However, the direct precursors of the two pyrazole nitrogen atoms and the biosynthetic machinery driving N-N bond formation have remained obscure.

To answer these long-standing questions, we sequenced the genome of the PZN producing strain *Streptomyces candidus* NRRL 3601, and identified a ~28 kb gene cluster as a candidate for the PZN biosynthetic gene cluster, which we name here as the *pyr* cluster (**Fig. 2a, Table S1**). This gene cluster carries a four-gene sub-cluster (*pyrOPQS*) that has been linked to the biosynthesis of coformycin, whose production seems to be associated with many nucleoside family natural products.^[17–19]^ Moreover, the *pyr* cluster also contains *pyrE*, which shows sequence homology to genes encoding β-ribofuranosylaminobenzene 5′-phosphate (β-RFAP) synthase. In the biosynthesis of methanopterin, β-RFAP synthase catalyzes the condensation of *p*-aminobenzoic acid with phosphoribosylpyrophosphate through a *C*-glycosidic linkage.^[20]^ A similar reaction for *C*-glycosidic bond formation seems to be also required during PZN biosynthesis. Furthermore, another interesting feature about the *pyr* cluster is the presence of *spb38* and *spb40* homolog genes *pyrM* and *pyrN*, indicating that this cluster encodes a potential hydrazine-producing pathway analogous to that of the s56-p1 pathway. which is also consistent with the N-N bond containing structure of PZN.^[7]^ We note that the *pyr* cluster share many homologous genes with the recently reported formycin biosynthetic gene cluster (*for* cluster).^[19]^ Considering the great structural similarity between pyrazomycin and formycin, it is not unexpected that they utilize similar biosynthetic genes. Taken together, our *in silico* analysis suggests that the *pyr* cluster might be responsible for PZN assembly in *S. candidus* NRRL 3601.

**Figure 2.**
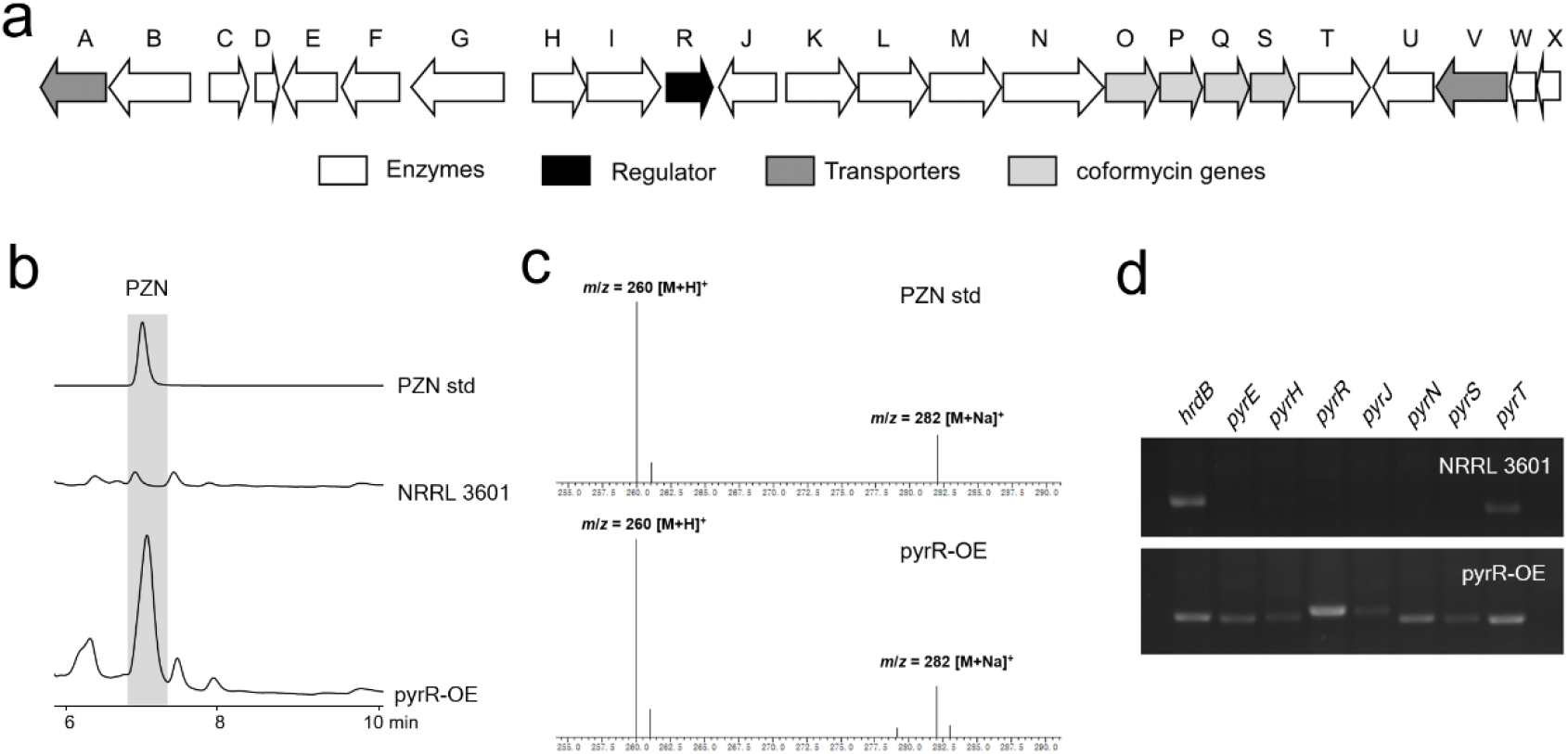
Identification of the gene cluster and cluster-situated activator for pyrazomycin biosynthesis. (**a**) The biosynthetic gene cluster of pyrazomycin. (**b**) HPLC analysis of the culture supernatants from strain *S. candidus* NRRL 3601 and its pyrR-overexpression mutant pyrR-OE. (**c**) MS analysis of the pyrazomycin standard compound and the new product produced by strain pyrR-OE. (**d**) Transcriptional analysis of selected genes from *pyr* gene cluster in strain NRRL 3601 and pyr-OE by RT-PCR.

Although strain *S. candidus* NRRL 3601 was known as a PZN producer, we failed to detect any PZN production with various media we tested, including the one used in early studies (**Fig. 2b–2d**).^[15]^ Introduction of a vector carrying the whole putative *pyr* cluster into the common *Streptomyces* host *S. albus* J1074 also did not produce any detectable PZN. To explore why no PZN is produced, we isolated total RNA from *S. candidus* NRRL 3601 and performed transcriptional analysis of the *pyr* cluster by RT-PCR. We found that most of the *pyr* genes were not actively transcribed under the culture condition we used (**Fig. 2d**). We thus turned to regulatory engineering of the PZN pathway. Analysis of the *pyr* genes revealed the presence of a single putative transcriptional regulator *pyrR*, which shows sequence homology to StrR-family regulators. As the prototype of this family, the streptomycin biosynthesis regulator StrR is the transcriptional activator controlling the expression of streptomycin biosynthetic genes by interacting with multiple promoter regions within the gene cluster.^[21,22]^ Members of this family also include NovG and Bbr, which are located in the biosynthetic gene clusters of novobiocin and balhimacin, respectively.^[23,24]^ This bioinformatic analysis suggested that *pyrR* likely encodes a pathway-specific activator of the PZN biosynthetic gene cluster. To interrogate whether *pyrR* could activate this biosynthetic pathway, we introduced an additional copy of *pyrR* into *S. candidus* NRRL 3601 under the control of a constitutive promoter to release the transcriptional control from higher-level regulatory mechanisms. Subsequent metabolic profiling of the resulting strain *S. candidus* pyrR-OE by LC-MS together with NMR analysis of isolated PZN demonstrated that PZN production was successfully recovered, with a PZN yield of ~10 mg/L (**Fig. 2b and 2c, Fig. S1**). Furthermore, RT-PCR analysis of selected *pyr* genes revealed their active transcription in *S. candidus* pyrR-OE, including the *spb40* homolog *pyrN*, which is located in a putative operon consisting of genes *pyrKLMN* (**Fig. 2d**). Altogether, these results demonstrated that PyrR is a transcriptional activator governing PZN biosynthesis, and by extension, the *pyr* cluster is responsible for PZN biosynthesis.

We next interrogated the biosynthetic origin of N-1 and N-2 in the pyrazole ring of PZN, which remain obscure despite the previous isotope feeding experiments and our *in silico* analysis of the *pyr* cluster. The presence of *spb38* and *spb40* homologs *pyrM* and *pyrN* in the *pyr* cluster indicate that a N-N bond forming mechanism analogous to that of the s56-p1 pathway might operate in PZN biosynthesis.^[7]^ We envisaged two possible routes that could afford the hydrazine moiety in the pyrazole ring (**Fig. 3a**). In one scenario, the Spb40 homolog PyrN could catalyze N-N bond formation between the *N*-6 nitrogen of l-*N*^6^-OH-Lys with another amine-containing substrate to generate a molecule as a hydrazine carrier, which would be followed by the transfer of the hydrazine moiety to a glutamate or α-KG derivative by downstream pathway enzymes. In this scenario, PyrN could be functionally equivalent to Spb40 and produce the product Lys-Gly for the subsequent hydrazine transfer. A second scenario involves the direct installation of hydrazine moiety on glutamate or its derivative, through linking the α-NH_2_ of glutamate (or its derivative) to N-6 nitrogen of l-N^6^-OH-Lys. The resulting product could then be processed by other *pyr* enzymes to furnish the pyrazole ring. To distinguish between these two possible routes, we first separately fed l-^15^N-Gly or l-^15^N-Glu (at concentrations of 3 mM) into the culture of *S. candidus* pyrR-OE and analyzed the isotope incorporation of PZN by LC-MS. We found that the nitrogen from the α-NH_2_ of glutamate efficiently incorporated into PZN, with the relative intensity of +1 Da peak increasing from 11% to 46% when compared to unlabeled PZN (**Fig. S2**). Furthermore, the +2 Da peak increased to 16 %. By contrast, only minor incorporation of the nitrogen from ^15^N-Gly was detected. Although it is likely that the observed incorporation pattern is due to or partially attributed to scrambling of the labels by primary metabolism, this result favors the second scenario, where the N-1, and by extension, C-3 to C-6 atoms, in the pyrazole moiety might derive from l-Glu as an intact unit (**Fig. 3a**).

**Figure 3.**
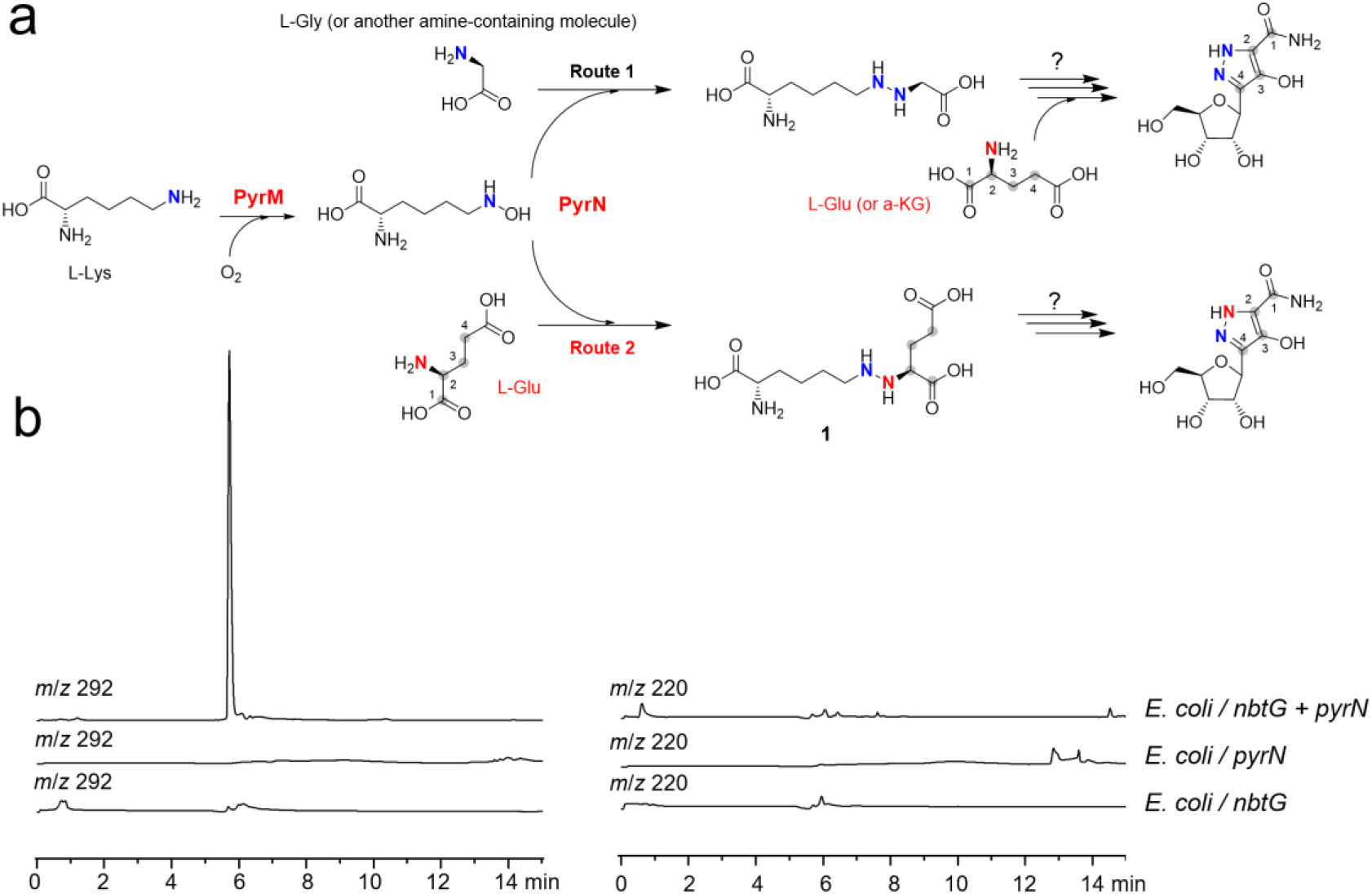
(**a**) Possible biosynthetic routes to the pyrazole ring in pyrazomycin based on previous studies. (**b**) LC-MS analysis of engineered *E. coli* strains carrying different gene(s).

To gain further insights into the product from the PyrN-catalyzed reaction, we adopted an *in vivo* reconstitution method by introducing *pyrN* into the *E. coli* strain expressing *nbtG*. NbtG, a homolog of PyrM and Spb38, is a well-characterized lysine *N*^6^-hydroxylase and thus could provide endogenous l-OH-*N*^6^-Lys *in vivo*. LC-MS analysis of culture supernatant from the *E. coli* strain carrying both *nbtG* and *pyrN* revealed the presence of a molecule with a MS signal at *m*/*z* 292, which is absent from that of *E. coli* strain expressing either *nbtG* or *pyrN* (**Fig. 3b**). This MS signal (*m*/*z* 292) is consistent with the [M+H]^+^ ion of a Lys-Glu conjugate product (**1**) (**Fig. 3a**). No MS signal (*m*/*z* 220) of Lys-Gly was detected, supporting that Gly is not a substrate of PyrN, in contrast to Spb40. To track the origin of compound **1**, we fed the strain *E. coli*/*nbtG*+*pyrN* with different combinations of ^13^C/^15^N-labeled and unlabeled amino acid precursors, including l-^13^C_6_-Lys, l-^15^N-Glu, l-^15^N-Gly (**Fig. S3**). Compared with cultures supplemented with unlabeled amino acids, the compound **1** produced in the presence of l-^13^C_6_-Lys showed the appearance of a strong +6 Da MS signal (*m*/*z* 298.1), supporting the incorporation of an intact lysine carbon skeleton. While **1** produced with the addition of ^15^*N*-Gly showed a similar isotope pattern to the one fed with unlabeled amino acids, addition of ^15^*N*-Glu resulted in significant enrichment of +1 Da signal, with an increase in the relative intensity from 12 % to 45 %. Fmoc-chloride treatment of the *E. coli* culture supernatants supplemented with ^13^C_6_-Lys, followed by LC-HR-MS analysis, resulted in the detection of a strain-specific metabolite from *E. coli*/*nbtG*+*pyrN* with MS signals at *m*/*z* 514.2164 and 520.2364, which are consistent with the [M+H]^+^ ion of Fmoc-Lys-Glu and Fmoc-(^13^C_6_)Lys-Glu conjugate, respectively (**Fig. 4a and Fig. S4**). Our attempt to isolate this molecule for NMR analysis was hampered by its low yield and instability. However, the fragmentation pattern of **1** from LC-HR-MS/MS analysis after Fmoc-Cl derivatization, together with the sequence homoloy between PyrN and Spd40, supports that **1** most likely arises from linking the α-NH_2_ of glutamate with the ε-NH_2_ of lysine (**Fig. 4b and 4c**).^[7]^ Altogether, the above results suggested that PyrM and PyrN together mediate the conjugation l-Glu and l-Lys though a N-N linkage.

**Figure 4.**
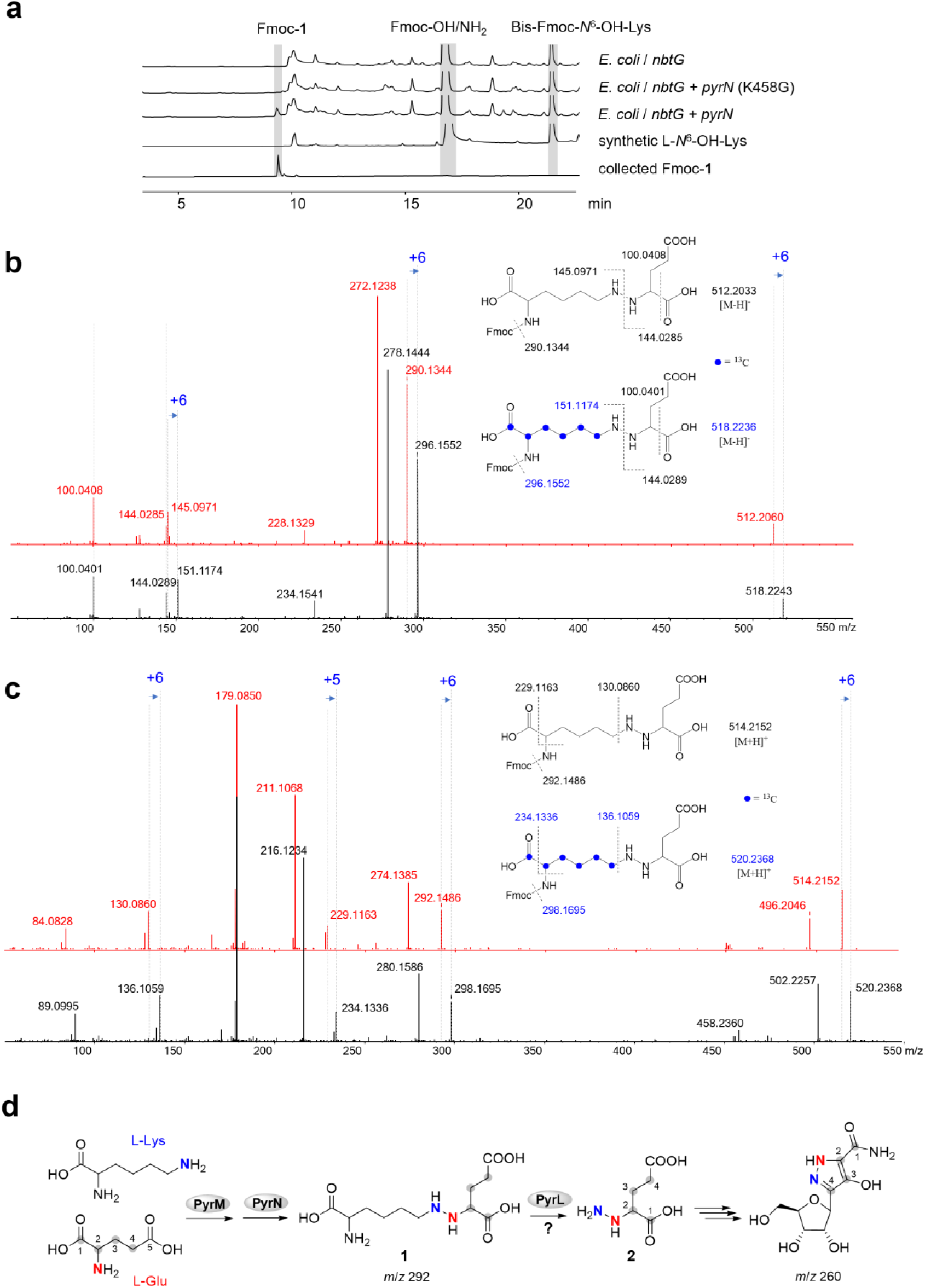
The *in vivo* reconstitution of the PyrN-catalyzed reaction in *E. coli* and proposed biosynthetic origin of the pyrazole ring in pyrazomycin. (**a**) HPLC analysis of culture supernatants from the *E. coli* strains containing different gene combinations after Fmoc-Cl derivatization. Note: Lys458 is a residue potentially involved in ATP-binding in PyrN (see below in the main text). (**b**) and (**c**) LC-HR-MS/MS anaylsis of the strain-specific metabolites from *E. coli* / *nbtG* + *pyrN* supplemented with l-^13^C_6_-Lys under negative and positive modes, respectively. Note: the red traces are from the MS/MS analysis of unlabeled molecules, and the black traces are from ^13^C_6_-labeld molecules. (**d**) Proposed biosynthetic origin of the pyrazole moiety in pyrazomycin based on the results from this study and previous stable-isotope precursor feeding experiments.

As in Spb40, the residues potentially involved in ATP-binding and metal-chelating are also conserved in the aaRS-like and cupin domains of PyrN, respectively (**Fig. S5 and S6**). Consistent with this *in silico* analysis, inductively coupled plasma mass spectrometry (ICP-MS) analysis of the purified His-tagged PyrN demonstrated that it contains two equivalents of zinc, supporting the presence of a single, conserved zinc coordination site in each domain (**Fig. S7**). Even though we were able to obtain soluble recombinant PyrN, our preliminary screening of reaction conditions for the *in vitro* assay of PyrN did not produce any detectable product so far, which could be attributed to yet-unknown factors necessary for this reaction. Nonetheless, we expect that the PyrN reaction mechanism is likely to be similar to that has been suggested for Spb40, where there is a similar lack of *in vitro* work reported in the literature.^[7]^ Based on the results from our *in vivo* reconstitution in *E. coli* and previous isotope tracking experiments on Spb40 (**Fig. S8**), the C-terminal aaRS-like domain of PyrN is likely responsible for glutamate carboxylate activation, to afford glutamyl-adenylate or further to give glutamyl-tRNA, which could then undergo ligation to the hydroxyl group of l-OH-*N*^6^-Lys to form an ester intermediate, and followed by rearrangement to generate **1**, which is likely mediated by the N-terminal cupin domain. Formation of an ester intermediate by the conjugation of an amino acid to a hydroxylamine, using aminoacyl-tRNA as a carrier, is not unprecedented. In the biosynthesis of azoxy-containing valanimycin, the seryl-tRNA synthetase VlmL produces l-seryl-tRNA, followed by VlmA-mediated seryl transfer to isobutylhydroxylamine to form O-seryl-isobutylhydroxylamine, which then undergoes subsequent transformation(s) for N-N bond formation.^[9]^ Following the formation of **1**, the next step in the PZN pathway is likely catalyzed by PyrL, whose encoding gene *pyrL* appears to be located in the same operon with *pyrM* and *pyrN*, indicating their functional relevance (**Fig. 2a**). PyrL displays sequence homology to saccharopine dehydrogenase. In lysine metabolism, saccharopine dehydrogenase mediates reversible conversion of saccharopine to l-2-aminoadipate-6-semialdehyde (AASA) and l-Glu (**Fig. S9**).^[25]^ PyrL might thus catalyze an analogous reaction, converting **1** to AASA and 2-hydrazinoglutaric acid (**2**), the latter of which could then be processed by downstream pathway enzymes for pyrazole ring assembly (**Fig. 4d**).

In conclusion, we have identified the gene cluster responsible for pyrazomycin biosynthesis in *S. candidus* NRRL 3601, and demonstrated that PyrR is a new member of the StrR family transcriptional activator controlling PZN biosynthesis. Furthermore, our current data strongly supports that PyrN is a new N-N bond forming enzyme, and the formation of the lysine-glutamate conjugate through a N-N linkage could be the starting point for the PZN biosynthetic pathway. This study paves the way for further biochemical characterization of PyrN and other PZN pathway enzymes involved in the assembly of the unique pyrazole ring.

## Experimental Section

Full experimental details are available in the Supporting Information

## Acknowledgements

This work was supported by funding from National Key R&D Program of China (2018YFA0903203), the National Natural Science Foundation of China (31872625), the Zhejiang Provincial Science Foundation (LR19C010001) and the Natural Sciences and Engineering Research Council of Canada (RGPIN-2016-03778).

