## Supplementary Information for "The biosynthetic gene cluster of the *C*-nucleoside antibiotic pyrazomycin with a rare pyrazole moiety"

**TABLE OF CONTENTS**

**Experimental Section**

**Supplementary Tables**

**Table S1.** The biosynthetic gene cluster of pyrazomycin.

**Table S2.** Primers used in this study**.**

**Supplementary Figures**

**Figure S1.** $^{1}$H- (top, 600 MHz) and $^{13}$C-NMR (bottom, 150 MHz) analysis of isolated PZN.

**Figure S2**. LC-MS analysis of PZN produced from strain *S. candidus* pyrR-OE fed with labeled precursors

**Figure S3**. LC-MS analysis of the incorporation pattern of compound (*m*/*z* 292)

**Figure S4**. LC-HR-MS analysis of the Fmoc-**1**

**Figure S5**. *in silico* analysis of the PyrN C-terminal aaRS-like domain

**Figure S6**. *in silico* analysis of the PyrN N-terminal cupin domain

**Figure S7**. SDS-PAGE and ICP-MS analysis of purified PyrN

**Figure S8**. Proposed PyrN-catalyzed reaction mechanism

**Figure S9**. Proposed PyrL-catalyzed reaction

**References**

### Experimental Section

##### General methods

DNA primers were purchased from Tsingke Biological Technology. Reagents were purchased from Sigma-Aldrich, New England BioLabs, Bio-Rad, Bio Basic Inc., and Sangon Biotech. DNA manipulations in *Escherichia coli* and *Streptomyces* strains were carried out according to standard procedures.[1,2] *Streptomyces* and its mutant strains were normally maintained on MS agar or ISP2 agar. Ampicillin (100 µg/ml), apramycin (50 µg/ml) and kanamycin (50 µg/ml), spectinomycin (50 µg/ml) were used for selection of recombinant *E. coli* and *Streptomyces* strains. Genome sequencing and *in silico* analysis of genome sequence was performed similarly as described previously,[3] with assembly of 9.13 megabase pairs of nonredundant sequences over 58 contigs with 252-fold read coverage. A genomic library of *Streptomyces candidus* NRRL 3601 was constructed as described previously, albeit with vector pOJ436.[3] Cosmid 14G6 containing the whole pyrazomycin biosynthetic gene cluster was observed by library screening with primers targeting selected *pyr* genes. Nucleotide sequence of the pyrazomycin gene cluster was deposited into the Genbank database under the accession number MN170532.

##### Construction of *S. candidus* pyrR-OE

The coding region of *pyrR* was amplified by PCR using NEB Phusion DNA polymerase and primers listed in **Table S2**, and inserted into NdeI/XbaI site of pYLD20 under the constitutive *ermE**p promoter.[4] After confirmation by DNA sequencing, the integrative plasmid pYLD20-*pyrR* was introduced into *S. candidus* NRRL 3601 by conjugation, to give the engineered strain *S. candidus* pyrR-OE. Four independent conjugates were used for subsequent metabolic analysis.

##### Metabolic analysis for the *S. candidus* strains

Spore suspensions of *S. candidus* strains were used to inoculate 250-mL flasks containing 50 mL of tryptic soy broth (BD) medium, which were incubated with shaking for 18 h at 30°C. 2.5 ml samples of the seed cultures were then used to inoculate 250 mL flasks containing 50 mL of the PROS medium [(g/L): soybean meal (15.0), casamino acid (1.0), $NaNO_3$ (3.0), glucose (20.0) and $CaCO_3$ (2.0)], and these were incubated with shaking at 30°C. In the case of isotope-labeled precursors feeding experiments, the isotope-labeled amino acids were added at final concentrations of 3 mM at 24 h. After 72 h, cultures were centrifugated at 12,000 rpm for 5 min to remove the mycelium. The supernatants were deproteinated through methanol precipitation, and followed by removal of methanol using SpeedVac. The resulting samples were then subjected to HPLC analysis. For isolation of pyrazomycin produced by *S. candidus* pyrR-OE, six flasks of 50 mL cultures were processed as described above, and then concentrated before fractionization on Sephadex LH-20 with MeOH:$H_2O$ (1:1) elution. The fractions containing the target molecule were combined, concentrated and subjected to preparative HPLC (YMC-Triart C18, 5 µm, 10 mm ID x 250 mm) for isolation of the pure compound. The $^1H$- and $^{13}C$-NMR spectra were recorded on a Bruker AV-600 MHz spectrometer using $D_2O$ as the solvent.

Analytic HPLC analysis was carried out with an Agilent 1260 HPLC apparatus using a Luna C18(2), 5 µm, 4.6 mm ID × 250 mm column (Phenomenex). Elution was performed at 0.6 mL/min with a mobile-phase mixture consisting of a linear gradient of water and acetonitrile ((v/v): 98:2, 0 to 10 min; 0:100, 10 to 20 min; 0:100, 20 to 25 min), both of which contain 0.05% (v/v) formic acid. A pyrazomycin standard compound (Sigma-Aldrich) was used as a control (detection wavelength: 254 nm). LC-MS was performed under the same conditions (except that the flow rate is 0.3 mL/min for LC-MS) on the Shimadzu LC/MS-

8045 system operated in positive mode.

**RNA isolation and RT-PCR**

The total RNA of *S. candidus* strains were isolated from the strain grown in PROS medium at 36 h. RNA was prepared with Trizol (Sangon) according to the manufacturer's instructions. Genomic DNA was removed by RNase-free DNase I (Takara), and the absence of genomic DNA contamination was confirmed by PCR using primer pairs targeting *hrdB*. The RNA integrity was checked by agarose gel electrophoresis, and its concentration was determined by using Nanodrop (Thermo Fisher Scientific). Two-step RT-PCR was performed; cDNA was first made from 1 μg total RNA using a Primescript cDNA Synthesis Kit (Takara) based on the manufacturer's manual, and cDNA was amplified by Taq DNA polymerase (Bio Basic, Inc.). The PCR conditions consisted of one cycle of denaturation at 94°C (2 min), followed by 30 cycles of 94°C (30 s), 58°C (30 s), and 72°C (60 s), with one extension cycle at 72°C (5 min).

***In vivo* reconstitution of the PyrN-catalyzed reaction in *E. coli* host**

Codon-optimized gene *nbtG* was synthetized and cloned into pCDFDuet-1 using NdeI and XhoI restriction sites to give pCDF-*nbtG*. Gene *pyrN* was amplified from cosmid 14G6 and cloned into pET28a using NdeI and XhoI restriction sites to afford pET28a-*pyrN*. For the construction of strain *E. coli* / *nbtG* + *pyrN*, pCDF-*nbtG* and pET28a-*pyrN* were co-transformed into *E. coli* BL21 (DE3). Similarly, strain *E. coli* / *nbtG* was constructed by co-transformation of pCDF-*nbtG* and pET28a empty vector. Strain *E. coli* / *pyrN* was constructed by co-transformation of pCDFDuet-1 empty vector and pET28a-*nbtG*. Strain *E. coli* / *nbtG* + *pyrN* (K458G) was constructed by co-transformation of pCDF-*nbtG* and pET28a-*pyrN* (K458G), in which the K458G mutation was introduced by using the Q5 Site-Directed Mutagenesis Kit (NEB) and confirmed by DNA sequencing. These *E. coli* strains were cultivated in M9 medium for protein expression and metabolites production. When the $OD_{600}$ reached ~0.6, the cultures were supplemented with IPTG at a final concentration of 0.1 mM, along with 0.05% (w/v) of amino acid precursors. The cells were then cultured at 30°C for another 12 h. The culture broth was centrifuged at 12,000 rpm for 5 min and the supernatant was collected and mixed with the 2 volume of acetonitrile. The resulting mixture was then centrifuged at 12,000 rpm for 5 min, and subjected to LC-MS analysis directly or after Fmoc-Cl derivatization, which was performed as described previously.[5] LC-MS was performed as described above. LC-HR-MS/MS was performed on waters UPLC coupled with an AB TripleTOF 5600+ mass spectrometer system using Waters ACQUITY UPLC BEH-C18 column (1.7 μm, 2.1 × 50 mm). The mobile phases were in water (A) and methonal (B), both of which contain 5 mmol ammonium formate. The linear gradient programs were as follows, 0/37, 2.5/100, 15/100 (min/B%), with a flow rate of 0.4 mL $min^{-1}$. MS/MS analyses were performed with a collision energy of 40 eV.

**Expression and purification of PyrN for ICP-MS analysis.**

The plasmid pET28a-*pyrN* was transformed into *E. coli* BL21 (DE3) for expression. To prepare starting culture, cells harboring pET28a-*pyrN* were grown overnight in Luria-Bertani (LB) broth with 50 μg/mL kanamycin at 37°C and 200 rpm. A starting culture (7.5 mL) was then used to inoculate LB broth (750 mL) containing 50 μg/mL kanamycin. The culture was grown at 37°C and 200 rpm to an optical density of 0.4 at 600 nm, and then cooled to 16 °C. Isopropyl β-D-1-thiogalactopyranoside (80 μg/mL) was added to induce overproduction of the protein. In one sample, 1 mM zinc acetate was also added. Following induction for 16 h, cells were harvested by centrifugation (4,300 x g, 15 min) and the pellet was resuspended in buffer (20 mM Tris pH 8.0, 500 mM sodium chloride, and 5 mM β-mercaptoethanol) containing 10 mM imidazole.

Next, cells were lysed through sonication (20% amplitude, 4 s on, 8 s off, 6 min) and separate by high-speed centrifugation (19,000 x g, 50 min). The clear lysate was gravity filtered through approximately 1 mL of Chelating Sepharose™ Fast Flow resin charged with nickel chloride. The protein was then eluted using a stepwise gradient of buffer containing increasing concentrations of imidazole (20 mM, 50 mM, 300 mM, and 500 mM) in 10 mL fractions. The fractions containing the protein were identified using SDS-PAGE, combined, and concentrated using Amicon® Ultra-15 Centrifugal Filter Units. The concentrated protein was then further purified via fast protein liquid chromatography in buffer (HiLoad™ 16/600 Superdex 200 pg column, flow rate 1 mL/min, monitored at 280 nm). Fractions containing the protein were again identified using SDS-PAGE and concentrated via spin centrifugation.

For inductively coupled plasma mass spectrometry, samples were diluted in buffer to 100 mM and sent to ALS Vancouver-Environmental in Burnaby, British Columbia, Canada. The purified proteins were diluted to 100 µM and treated with hydrochloric acid and nitrate acid to release all metal ions before testing.

### Supplementary Tables

**Table S1**. The biosynthetic gene cluster of pyrazomycin (GenBank accession number: MN170532)

| Genes | Size (aa) | Putative function |
|---|---|---|
| *pyrA* | 391 | MFS transporter |
| *pyrB* | 476 | Adenylosuccinate lyase |
| *pyrC* | 224 | HAD family hydrolase |
| *pyrD* | 123 | Cupin |
| *pyrE* | 334 | β-RFAP synthase |
| *pyrF* | 345 | SAICAR synthetase |
| *pyrG* | 540 | Fumarate reductase |
| *pyrH* | 310 | oxidoreductase |
| *pyrI* | 444 | monooxygenase |
| *pyrR* | 279 | StrR-like regulator |
| *pyrJ* | 351 | FAD dependent oxidoreductase |
| *pyrK* | 417 | Phosphoribosylglycinamide synthetase |
| *pyrL* | 400 | Saccharopine dehydrogenase |
| *pyrM* | 437 | L-Lysine 6-monooxygenase |
| *pyrN* | 668 | Cupin-Methionine-tRNA ligase |
| *pyrO* | 250 | SAICAR synthetase |
| *pyrP* | 233 | Short-chain dehydrogenase |
| *pyrQ* | 260 | HAD family phosphatases |
| *pyrS* | 284 | ATP phosphoribosyltransferase |
| *pyrT* | 407 | Phthalate 4,5-dioxygenase |
| *pyrU* | 387 | Amidohydrolase 3 |
| *pyrV* | 395 | MFS transporter |
| *pyrW* | 156 | NADH:FMN oxidoreductase |
| *pyrX* | 169 | NADPH-dependent FMN reductase |

**Table S2** Primers used in this study

| Primers | Sequence (5'→3') | Description |
|---|---|---|
| pyrR-NdeI-F | AGCAGCCATATGCAGTTGCTTCGTGAGGAAAGG | Primers for *pyrR* cloning |
| pyrR-XbaI-F | AGCAGCTCTAGACTAGTCCGACTTCTCGTCGAGTTC | |
| hrdB-F | CCAAGAACCACCTGCTCGAGGC | Primers for RT-PCR of *hrdB* |
| hrdB-R | GTCGAGGTAGTCCCGCAGAACC | |
| pyrE-F | GGAGCAGAACGTGTCGTAGTCG | Primers for RT-PCR of *pyrE* |
| pyrE-R | CTCAGCTTCACCCTGATCAGCC | |
| pyrH-F | CGACTACCTGGTGCTGCCTGAC | Primers for RT-PCR of *pyrH* |
| pyrH-R | GAGTCGTGAACCGCACCAGGAC | |
| pyrR-F | TGTCACCGTGCCGATCTCGTTG | Primers for RT-PCR of *pyrR* |
| pyrR-R | CGACTTCTCGTCGAGTTCGTCC | |
| pyrJ-F | TGAGCGTGTGGTCGAGGTCGTG | Primers for RT-PCR of *pyrJ* |
| pyrJ-R | TGAAGGTCCTGGTGATCGGTGC | |
| pyrN-F | GCACAACCACCACGACCTGGAG | Primers for RT-PCR of *pyrN* |
| pyrN-R | GACGAACGCTTCGAGCAGGAAC | |
| pyrS-F | GGTGTTCGGCAAACCTCGCGAG | Primers for RT-PCR of *pyrS* |
| pyrS-R | GAAGCTCGTAGGTCGCTTGTGC | |
| pyrT-F | GCTGCTGCGCGAGTACTGGATG | Primers for RT-PCR of *pyrT* |
| pyrT-R | AGGCTGTCCCCAGTGGACAGTC | |
| nbtG-NdeI-F | AGCAGCCATATGGAAACCCTGCTGGTTGTTG | Primers for *nbtG* cloning |
| nbtG-XhoI-R | AGCAGCCTCGAGTCACTGTGCTGCCAGCTGACGT | |
| pyrN-NdeI-F | AGCAGCCATATGATCGTCAGTGAGATCCCCC | Primers for *pyrN* cloning |
| pyrN-XhoI-R | AGCAGCCTCGAGTGTCGGTGTCGGTGTCGGTGT | |
| pyrN-K458G-F | GGCTTCTCCACCGGCCGCAAGCAC | Primers for point mutation of PyrN |
| pyrN-K458G-R | CTCCCCGTCGAGGAGGTAGAAG | |

### Supplementary Figures

**Figure S1.** [1]H- (top, 600 MHz) and [13]C-NMR (bottom, 150 MHz) analysis of PZN isolated from *S. candidus* pyrR-OE.

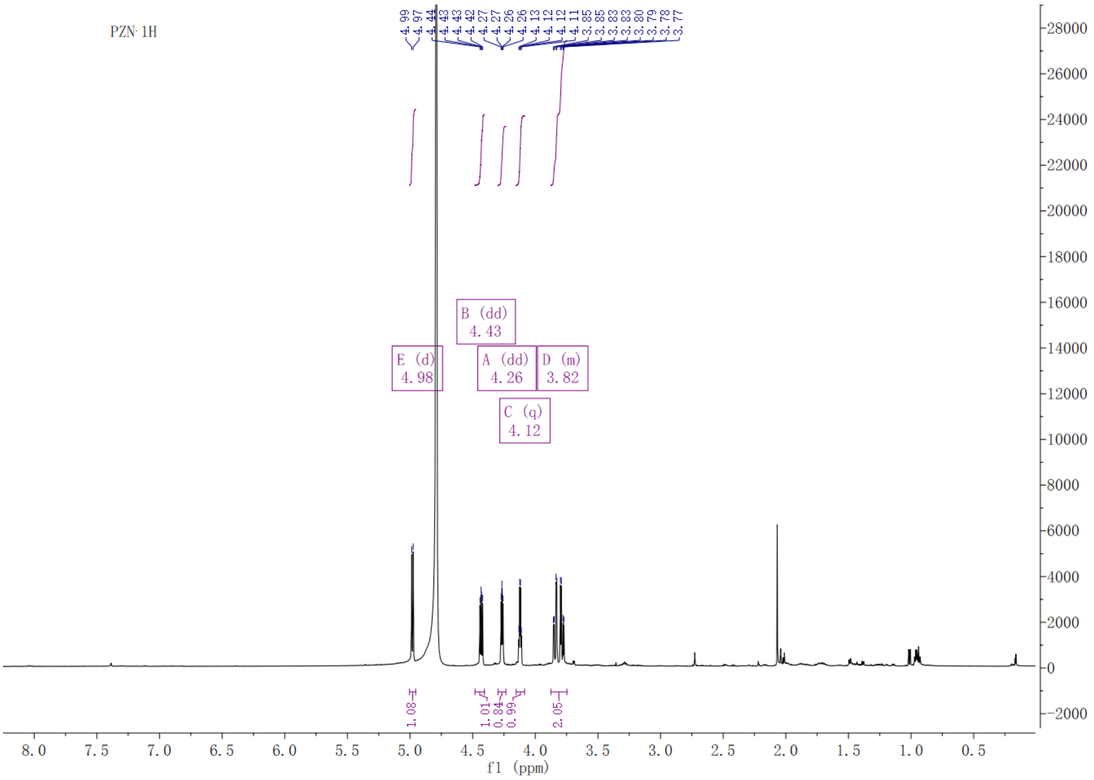


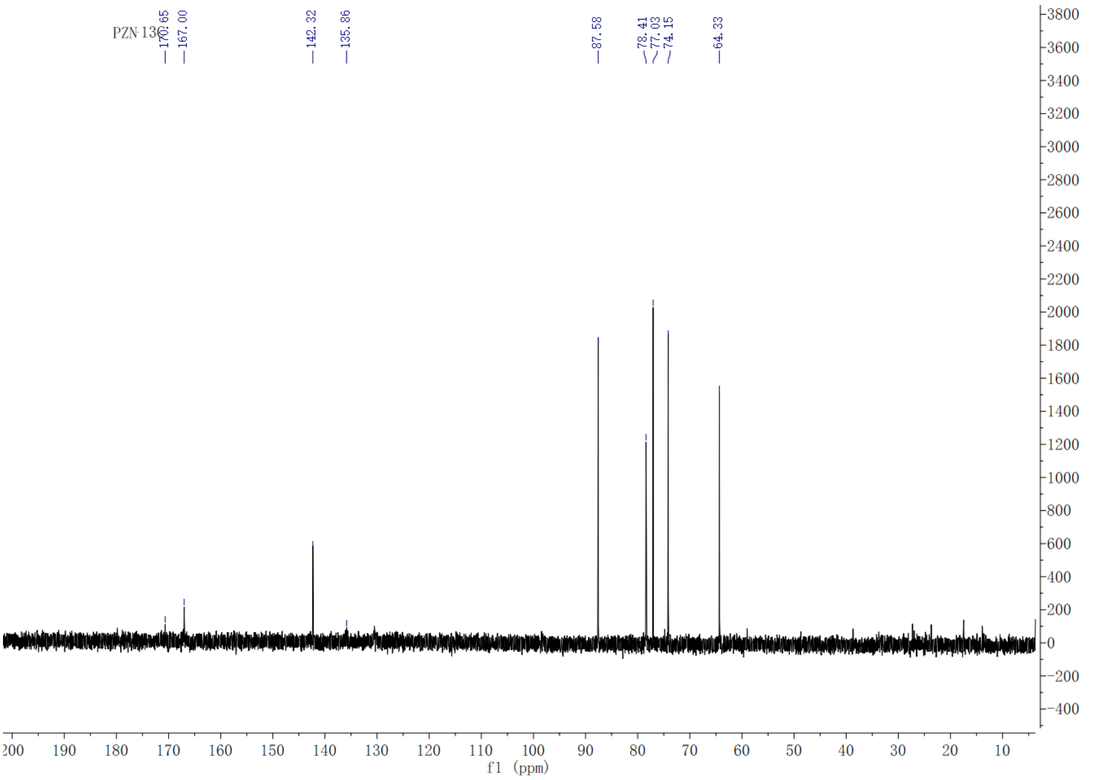

**Figure S2**. LC-MS analysis of PZN produced in the culture supernatants of strain *S. candidus* pyrR-OE supplemented with L-$^{15}$N-Glu or L-$^{15}$N-Gly. Relative intensity of the MS signals is indicated in parentheses.

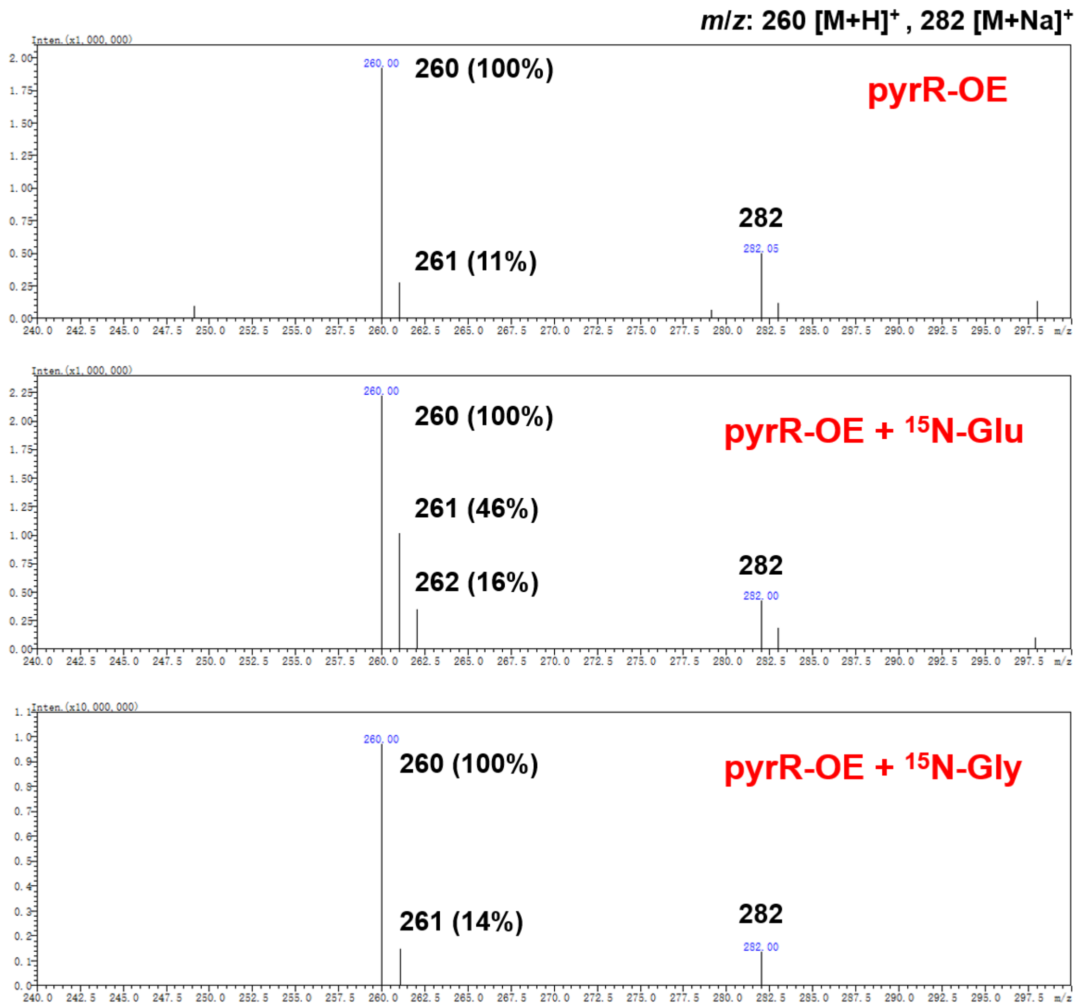

**Figure S3** Investigation of the incorporation pattern of compound (*m*/*z* 292) produced in the culture supernatants of *E. coli* / *nbtG* + *pyrN* fed with different precursors. Relative intensity of the MS signals are indicated in parentheses.

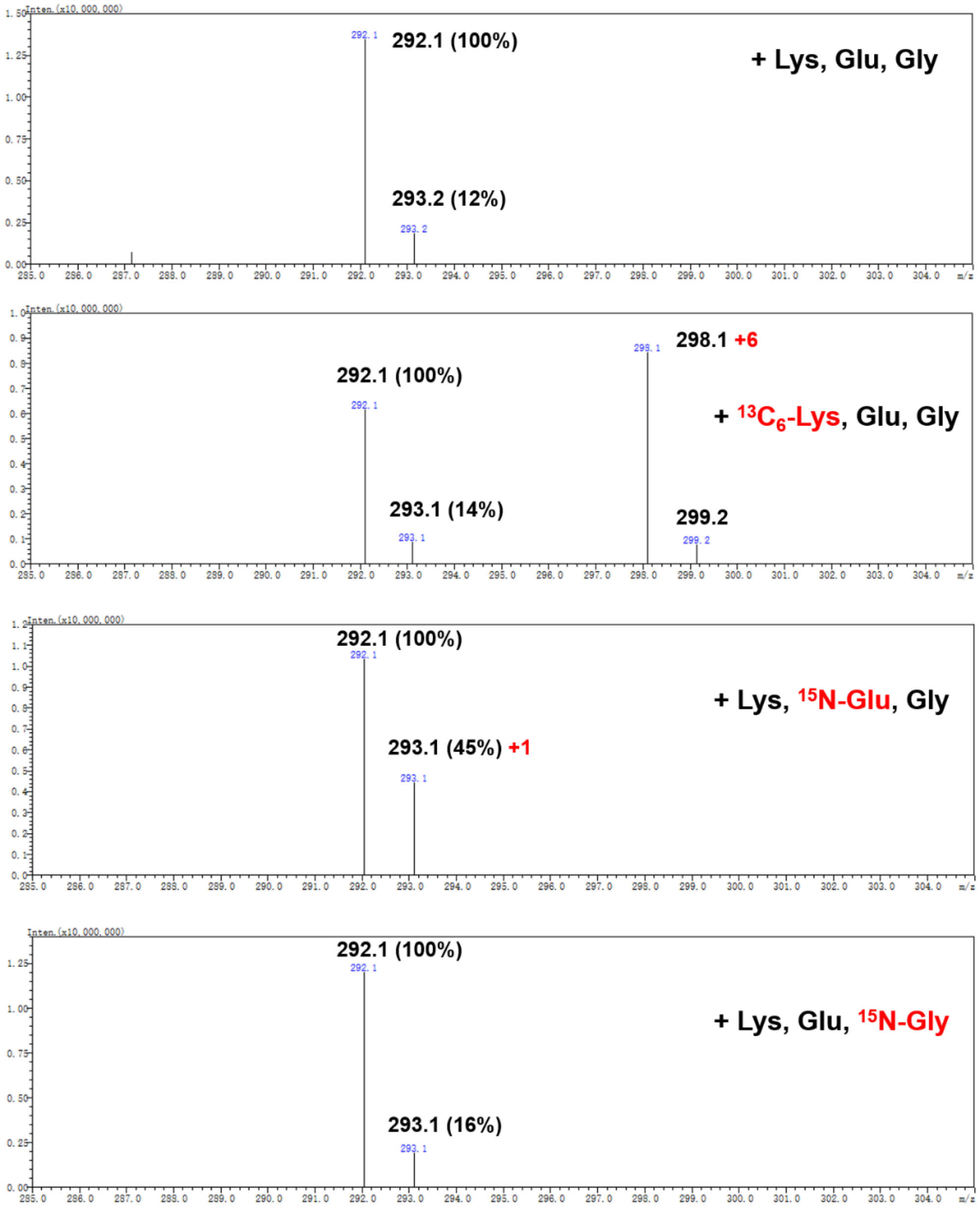

**Figure S4.** LC-HR-MS analysis of the Fmoc-derivatives of strain-specific metabolites from *E. coli* / *nbtG* + *pyrN* supplemented with L-$^{13}C_6$-Lys.

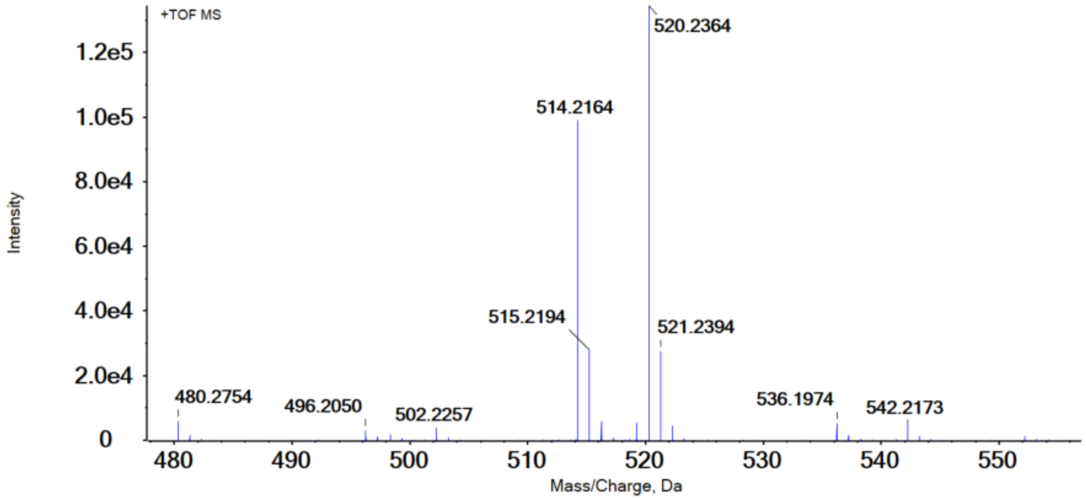

**Figure S5.** Sequence alignment (top) and homology model (bottom) of the C-terminal aaRS-like domain of PyrN based on template (PDB: 1F4L), which is a *E. coli* MetRS. Putative ATP-binding motifs are highlighted by purple boxes. Four conserved cysteine residues involved in zinc coordination are indicated with orange triangles. The lysine chosen for site-directed mutagenesis (Fig. 4a in main text) is indicated with a purple triangle. The above residues are also highlighted in the protein model structure.

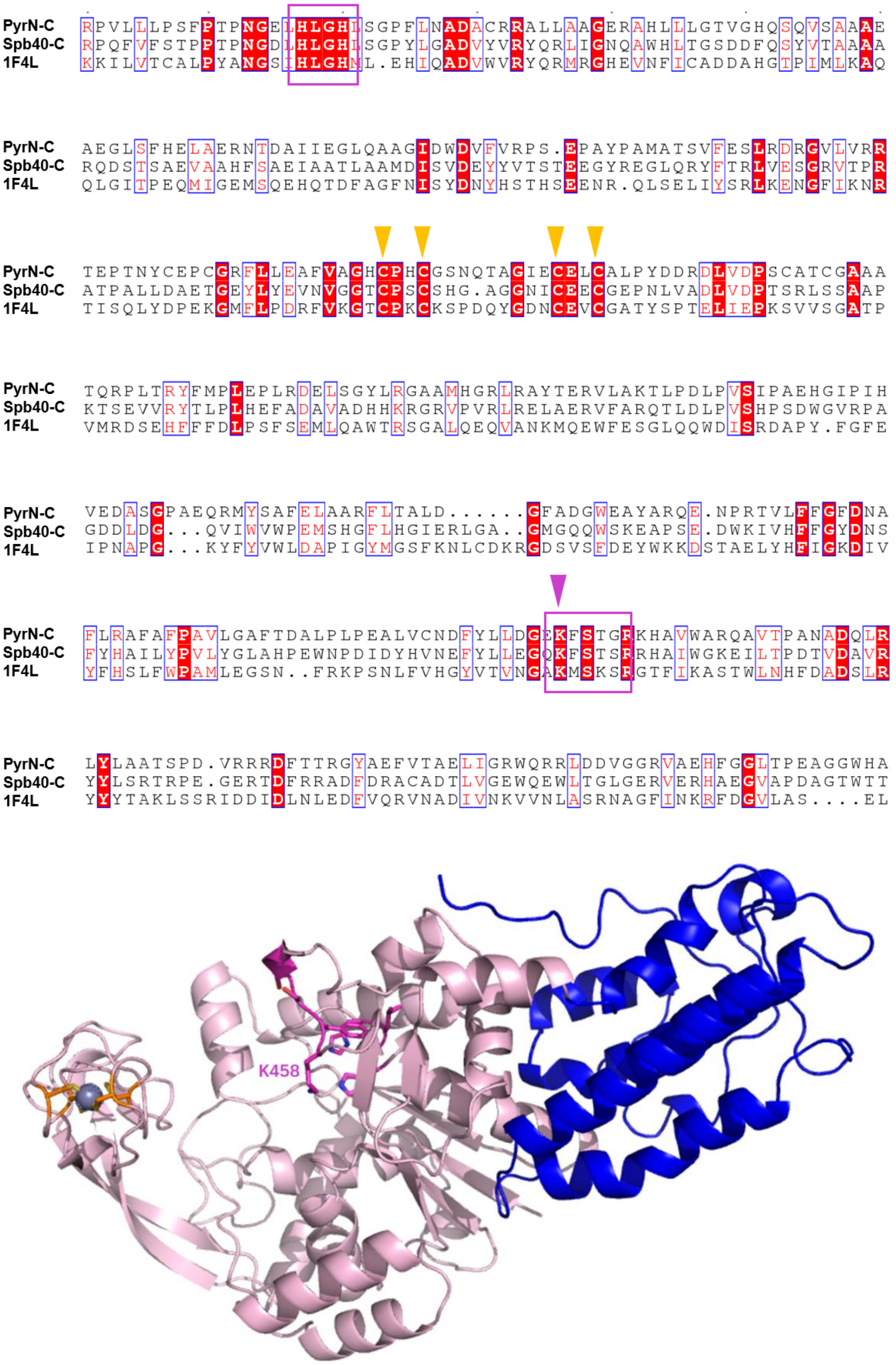

**Figure S6.** Sequence alignment (top) and superposition of the homology model of PyrN N-terminal cupin domain with the template (PDB: 5UQP), which is a zinc-binding cupin protein from *Rhodococcus jostii* RHA1, with unknown function. Residues potentially involved in metal coordination are indicated with orange triangles and indicated as sticks in the protein model structure.

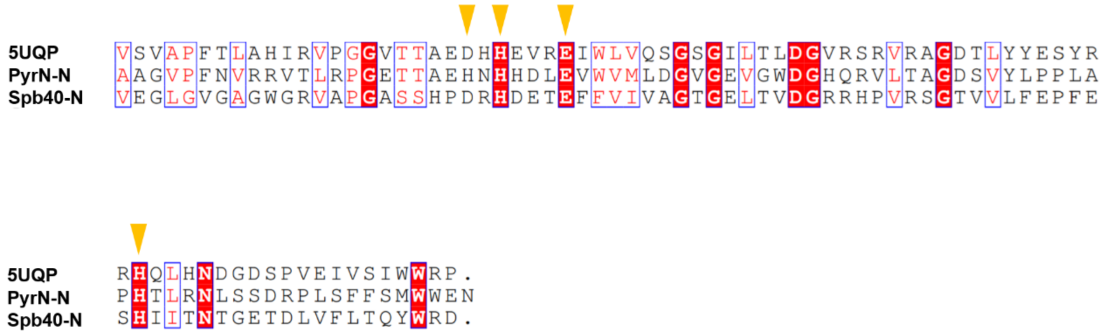


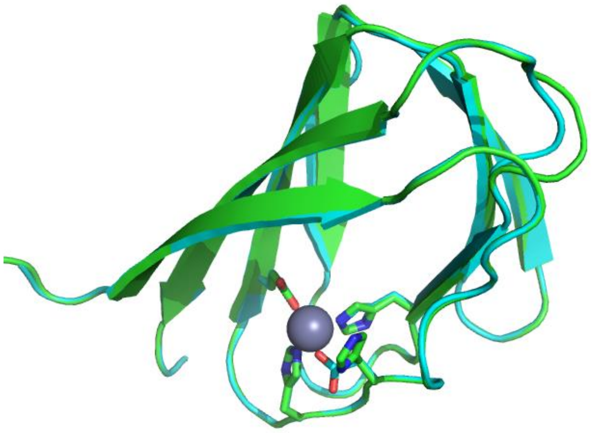

**Figure S7.** Inductively coupled plasma mass spectrometry (ICP-MS) analysis of purified 6×His-tagged PyrN. (A) SDS-PAGE analysis of N-terminal His-tagged PyrN. (B) ICP-MS analysis of purified PyrN. Data in gray indicates values below the detection limit. Note: the purified protein from un-supplemented LB broth contained 1.77 equivalents of zinc, accompanied by 0.43 equivalents of copper, and trace amounts of other metals. However, when the growth media was supplemented with 1 mM zinc acetate at the time of protein expression, the amount of zinc present increases to 2.01 equivalents while the amount of copper present drops below the detection limit. Therefore, in the latter case, the protein likely took up additional copper to compensate for the lack of available zinc. In both samples, the amount of nickel present is consistent with expected values of residual nickel bound to the His-tag. The presence of 2 equivalents of zinc supports predictions in S6 and S7, indicating the presence of a single, conserved zinc coordination site in each domain.

**(A)**

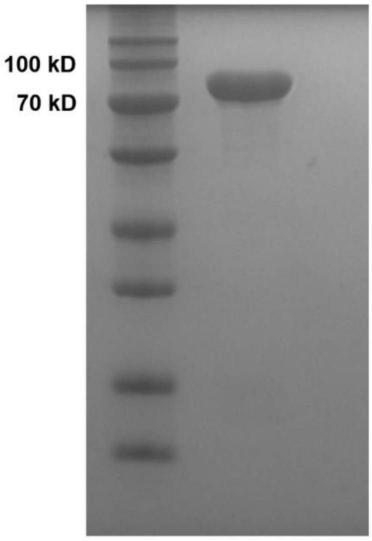


(B)

| | **PyrN (from LB media)** | | **PyrN (from LB media + 1 mM zinc acetate)** | |
|---|---|---|---|---|
| Metal | Observed concentration (μM) | Metal (mol): PyrN (mol) | Observed concentration (μM) | Metal (mol): PyrN (mol) |
| Co | < 0.034 | < $3.4 \times 10^{-4}$ | < 0.034 | < $3.4 \times 10^{-4}$ |
| Cu | 43 | 0.43 | < 9.2 | < 0.092 |
| Fe | 8.2 | 0.082 | < 3.6 | < 0.036 |
| Mg | 9.87 | 0.0987 | 4.11 | 0.0411 |
| Mn | 0.0983 | $9.83 \times 10^{-4}$ | 0.0819 | $8.19 \times 10^{-4}$ |
| Mo | < 0.010 | < $1.0 \times 10^{-4}$ | < 0.010 | < $1.0 \times 10^{-4}$ |
| Ni | 53.2 | 0.532 | 73.4 | 0.734 |
| **Zn** | 177 | **1.77** | 201 | **2.01** |

**Figure S8.** Proposed PyrN-catalyzed reaction involving formation and rearrangement of an ester intermediate.

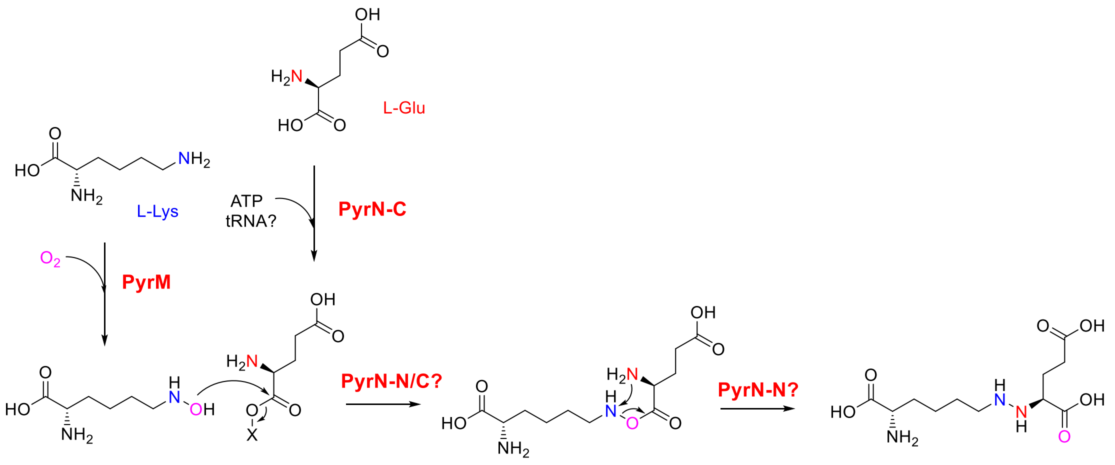

**Figure S9.** Saccharopine dehydrogenase-catalyzed reaction and the proposed PyrL-catalyzed reaction for the conversion of **1** to **2**.

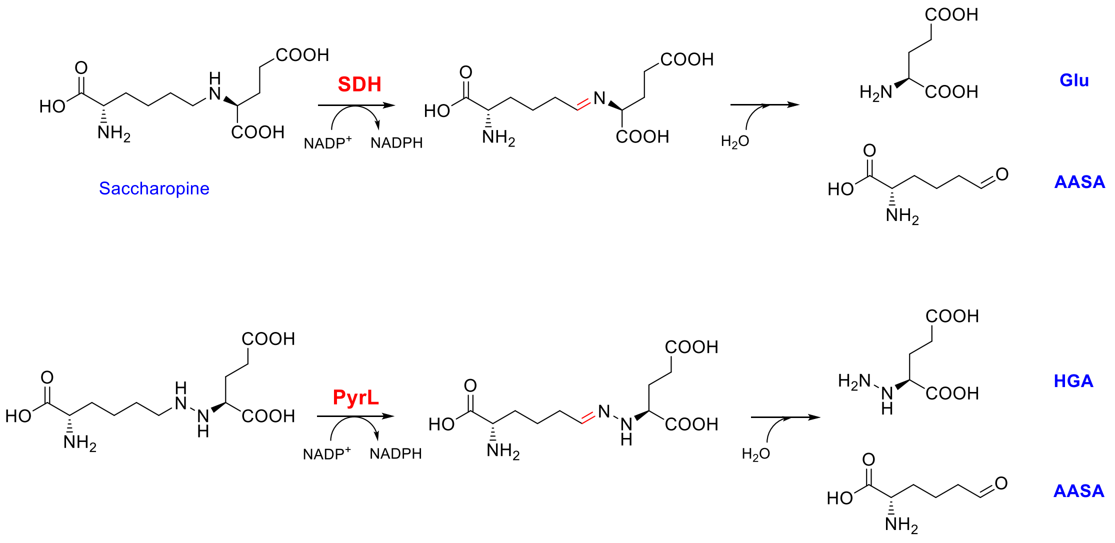
